# Rapid genomic expansion and purging associated with habitat transitions in a clade of beach crustaceans (Haustoriidae: Amphipoda)

**DOI:** 10.1101/2020.08.26.268714

**Authors:** Zachary B. Hancock, Faith O. Hardin, Archana Murthy, Andrew Hillhouse, J. Spencer Johnston

## Abstract

Genome sizes vary by orders of magnitude across the Tree of Life and lack any correlation with organismal complexity. Some crustacean orders, such as amphipods, have genome sizes that correlate with body size, temperature, and water depth, indicating that natural selection may constrain genome sizes due to physiological pressures. In this study, we examine the relationship between genome size, repetitive content, and environmental variables on a clade of sand-burrowing amphipods (Haustoriidae) that are distributed across the Gulf of Mexico and the North Atlantic. We uncover a 6-fold genome size variation within a clade that is less than 7 million years old. Unlike previous studies, we find no correlation between genome size and latitude, but do uncover a significant relationship between genome size and body length. Further, we find that the proportion of repetitive content predicts genome size, and that the largest genomes appear to be driven by expansions of LINE elements. Finally, we find evidence of genomic purging and body size reduction in two lineages that have independently colonized warm brackish waters, possibly indicating a strong physiological constraint of transitioning from surf-swept beaches to protected bays.

**Significance Statement:** The evolution of genome size has been a long-standing puzzle in biology. In this work, we find that genome sizes may be driven by different selection regimes following shifts to a new habitat. Dramatic genome size changes can occur rapidly, in only a few million years.

**Data Availability Statement:** Raw data sheets have been deposited on Dryad: SUBMITTED. Raw sequence reads are available at from NCBI under Bioproject SUBMITTED.

## Introduction

Genome sizes vary by several orders of magnitude across the Tree of Life and lack any correlation to organismal complexity (known as the *C*-value paradox; Gregory 2005; Lynch 2007). For example, the fern *Tmesipteris obliqua* has a genome fifty times larger than ours (Hidalgo et al 2017). Several theories have been proposed to explain this variation. The mutational-hazard hypothesis posits that genome size expansion is largely the result of the proliferation of parasitic DNA, which is made possible by relaxed selection on genomes (Lynch & Conery 2003; Lynch 2007; Lefébure et al. 2017). Therefore, in organisms with low effective population sizes (*N_e_*), we would expect a general increase in genome size due to weakened selection against these mildly deleterious expansions. In addition, large genomes are metabolically costly to replicate, slow the rate of replication and cell division (Kozlowski et al 2003; Gregory 2005) and increase cell size (Cavalier-Smith 1978), which may result in larger body sizes, especially within crustaceans (Jeffrey et al. 2016; Hultgren et al. 2018).

While genome size variation exists at large phylogenetic scales, in animals (but not plants, e.g. Hloušková et al 2019) it is largely constrained at smaller scales with organisms closely related having similar genome sizes (e.g., Tiersch & Wachtel 1991; Sessegolo et al 2016; Roebuck 2017; but see Lower et al 2017). This phylogenetic constraint has been used as evidence against the mutational-hazard hypothesis (Whitney & Garland 2010; but see Lynch 2011). Alternatively, congruence between genome size and phylogenetic relatedness could be a product of the lag in evolutionary timescales over which transposable elements (TEs) noticeably proliferate compared to reductions in long-term *N_e_*. Therefore a step toward disentangling the evolutionary rate of TEs and long-term *N_e_* variation on phylogenetic genome size congruence would be to identify young clades of organisms with dramatic differences in genome size that may represent a kind of “upper-bound” to TE expansion rates in nature.

Spatial variation in selection may also explain genome size differences across a clade. Since genome sizes impact metabolic rates, environmental factors that directly impact physiology may constrain genome size. Alfsnes et al. (2017), in a study across Arthropoda, found that for Hexapoda there was a strong phylogenetic signal in genome size that appeared to be related to developmental strategies, whereas in crustaceans the relationship was less reliant on phylogeny and more a product of maximum latitude and water depth. Hultgren et al. (2018) found correlations between genome size and latitude in Amphipoda, and Jeffrey et al. (2016) found that genome sizes of Lake Baikal amphipods were correlated with water depth, body size, and diversification rates. In addition, amphipods may show genome size variation within a cryptic species complex (Vergilino et al 2012).

Crustaceans, and many ectotherms in general, often follow a “temperature-size rule” (Walters & Hassell 2005) in which body size increases with decreasing temperature, whether at deeper waters or higher latitudes. Given that previous work has found an association between genome size, body size, and latitude in some crustacean groups (e.g., Hultgren et al 2018; Jeffrey et al. 2016; Alfsnes et al. 2017), it’s conceivable that there may be a general “genome-temperature-size rule”. Specifically, when body size is dictated by temperature, genome size should also be driven by temperature. This line of reasoning originates from the fact that at lower temperatures selection on metabolic rates is weaker, and therefore genome sizes are free to expand (Gregory 2005). Genome size expansion may reduce developmental rates and therefore generation times (Gregory & Johnston 2008), which may lead to an increase in body size.

Haustoriid amphipods are a family of beach-dwelling crustaceans that are widely distributed across the northern hemisphere. These amphipods are morphologically specialized to a fossorial lifestyle and display dramatic body size variation across their range (LeCroy 2002; Hancock & Wicksten 2018). In the Gulf of Mexico (GoM), haustoriids show an increase in body size west-to-east, with the largest in Florida and the smallest in Texas and Louisiana. A similar pattern of body size variation is seen in the Atlantic; however, body sizes increase south-to-north, with the smallest in Florida relative to the largest in Massachusetts. Therefore, haustoriid amphipods represent a natural system to test a possible “genome-temperature-size rule”. Since mean annual temperatures do not vary across the GoM but body sizes do, we do not expect to see a change in genome sizes if temperature is the constraining factor. Alternatively, in the Atlantic where mean annual temperatures vary dramatically from the southern to northernmost latitudes, we expect genome sizes to increase with increasing latitude. In this way, we can tease apart the correlation between body size and temperature seen in most amphipods (and ectotherms in general), and isolate the causal factor influencing genome size in this clade.

Previous work on this family has found strong population structure and widespread cryptic diversity (Hancock et al. 2019). In the GoM there are at least 6 species of haustoriids: *Haustorius galvezi, H. jayneae*, and a species complex of four distinct lineages of *Lepidactylus triarticulatus*. In the Atlantic, there exists only one “species” of *Haustorius, H. canadensis*, which ranges from at least Melbourne, Florida to Cape Cod, Massachusetts. This species shows incredible body size variation – at its lower range it is 4–5 mm in length, whereas at its northernmost range it can be as large as 18 mm (LeCroy 2002). Given the extent of cryptic diversity in this group already identified in the GoM (Hancock et al 2019), we expect that similar diversity exists in the Atlantic and that *H. canadensis* likely represents a cryptic species complex. Other genera exist in the Atlantic as well, but only *L. dysticus* shows a similar body size variation.

In this study, we evaluate genome size variation within haustoriid amphipods and test for correlations between environmental and phenotypic variables such as latitude, temperature, salinity, and body size to evaluate the existence of a general “genome-temperature-size rule”. We provide a time-calibrated phylogeny of the Haustoriidae to determine the timescale over which genome size variation occurs, and use this phylogeny to reconstruct ancestral environments, body sizes, and genome sizes. Finally, we perform low-coverage next-generation sequencing to characterize repetitive content across the genome to identify which families of repeat elements might be driving genome expansion.

## Materials & Methods

### Specimen collections and measurements

Amphipods were collected across 9 sites in the GoM covering roughly 20° longitude and 6 sites in the Atlantic spanning 20° latitude (**Figure 1a**). Specimens were collected in the swash zone using a hand-shovel and 435*μ*m sieve plate. Individuals targeted for genome size estimation were removed from the sieve plate with forceps and either kept alive in sand and seawater from their local environment or were fresh frozen on dry ice. For specimens in which genome sequencing was to be performed, they were removed from the plate and placed immediately into 95% EtOH. Specimens were identified to species in the lab using a Leica M205 FA dissecting scope, and we recorded body length for each specimen as distance (in mm) from rostrum to epimeron 3. At each site, we recorded both water temperature and salinity (using a refractometer). In addition, we categorized sand grain size at each location as “fine”, “medium”, and “course” based on how freely it passed through sieve plates of 1.18mm, 0.6mm, and 0.3mm. Sand denoted “fine” passed through all plates without retaining any shells or rocks; “medium” sand passed through the first two without retaining shells or rocks but not the final; “coarse” sand passed through the first and second, but large amounts of rocks and shell hash were retained.

**Figure 1.**
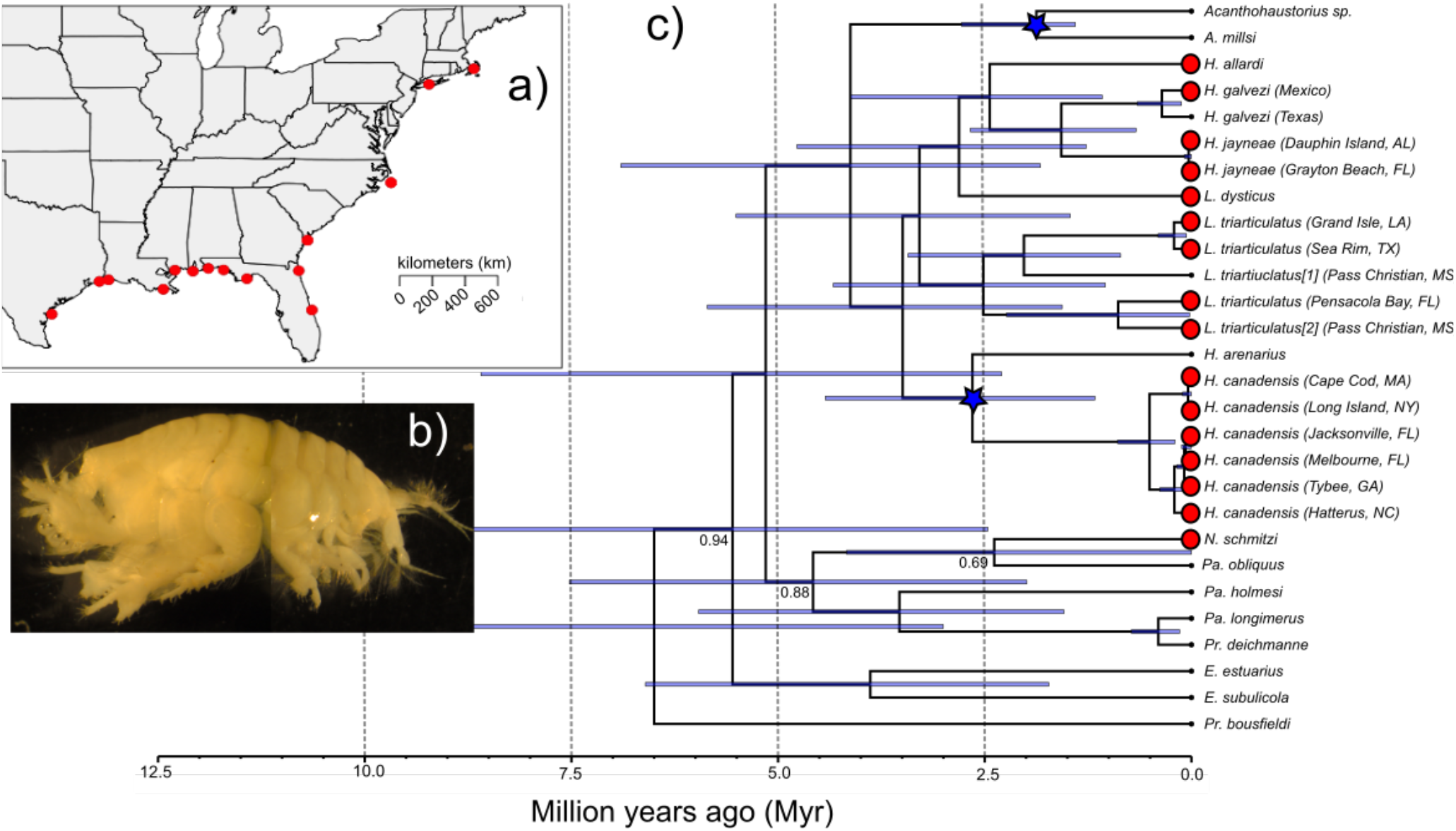
Calibrated phylogeny of the Haustoriidae. a) Map of sample sites; b) *L. triarticulatus* from Pensacola Bay, FL; c) time-calibrated phylogeny. Red circles are samples with genome size estimates; blue stars are calibration points; node bars represent 95% HPD. All nodes have PP > 0.95 unless specified.

### Genome size estimation

Genome size estimation via flow cytometry was performed on fresh frozen or live samples as described in Johnston et al. (2019) and Harahan & Johnston (2011). In brief, the anterior portion (excluding reproductive tissues) of each amphipod was placed into 1ml of Galbraith buffer in a 2ml Dounce tube (Kontes) along with the brain tissue from two standards, a female *Callosobrucus maculatus* (1C = 1240 Mbp) and a male *Periplaneta americana* (1C = 3338 Mbp). Nuclei from the sample and standard were released with 15 strokes of the loose (B) pestle. Debris was reduced by filtering the ground solution through a 40U nylon filter. DNA was stained for 2 hours in the cold and dark with prodidium iodide (25 ppm). Mean stain uptake in the 2C nuclei of the standards and the sample were determined using a CytoFlex Flow Cytometer (Beckman Dickenson). The genome size of each sample was calculated as the ratio of the mean fluorescence (output as a channel number) of the sample and the standard times the amount of DNA in the standard. Preliminary estimates found no sex-specific differences in genome size, and thus we pooled males and females. Unfortunately, sample degradation reduced our sample size to 1 for many sites, precluding us from evaluating within-site variation (**Table 1**).

**Table 1.**
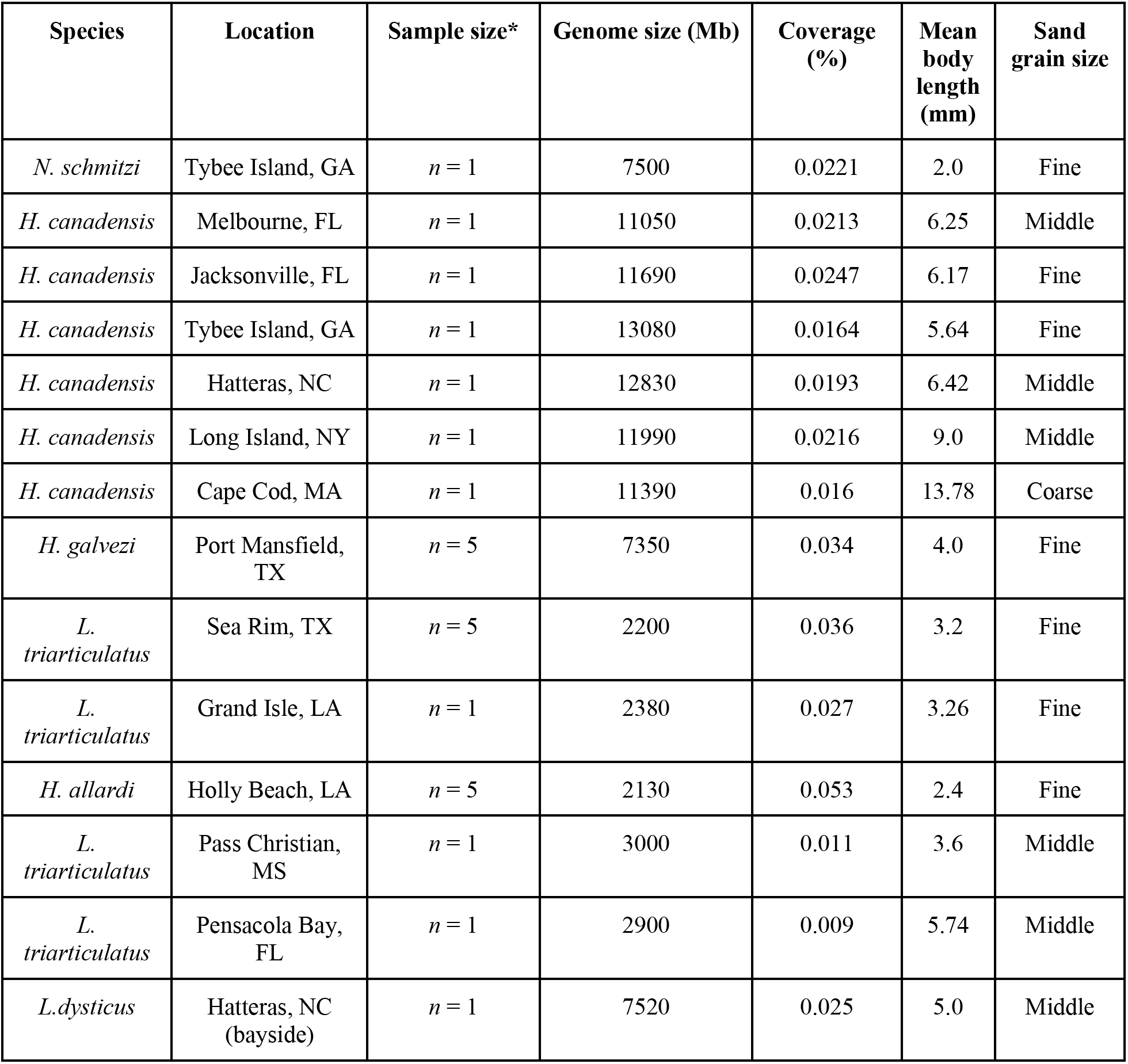
Summary of samples and their genomes sizes.

### DNA extraction and sequencing

Whole genomic DNA was extracted from either the whole specimen or pereopods 6–7 depending on the size of the amphipod using an EZNA Tissue DNA kit (Omega Bio-tek Inc.) following manufacturer’s protocols. Mitochondrial cytochrome oxidase I (*COI*) was amplified using forward and reverse primers from Folmer et al. (1994), and nuclear 28S ribosomal RNA (*28S*) was amplified using primers from Hancock et al. (2019). Polymerase chain reaction (PCR) conditions followed Hancock et al. (2019) for both mitochondrial and nuclear loci. Amplicons were verified using gel electrophoresis and purified with ExoSAP-IT (Affymetrix Inc.). Sanger sequencing on forward and reverse strands was performed at the DNA Analysis Facility on Science Hill at Yale University. Sequences were cleaned and manually edited using Sequencher v4.10.1 (Gene Codes Corp.), and alignments were performed in MAFFT 7 (Katoh & Standley 2013). The *COI* sequence was visually checked in Mesquite v3.5 (Maddison & Maddison 2018) to ensure there were no premature stop codons.

To generate repeat profiles, we performed low-coverage Illumina sequencing on 85 individuals from 15 locations across the GoM and North Atlantic (**Table 1; Figure 1a**). DNA was quantified with the dsDNA high sensitivity Qubit Assay (ThermoFisher) and checked for quality with an Agilent TapeStation genomic DNA tape (Agilent Technologies). Sequencing libraries were generated using the Swift 2S Turbo DNA Library preparation kit with enzymatic fragmentation and combinatorial dual indexing following the manufacturer’s instructions (Swift Biosciences). Libraries were pooled in equimolar ratios and sequenced in a single lane of 2 x150 paired-end sequencing on an Illumina NovaSeq 6000 (Illumina).

### Characterization of repeat elements

Quality control was performed on FASTQ files in the RepeatExplorer Galaxy platform (https://repeatexplorer-elixir.cerit-sc.cz/). For each population, individual left-hand and right-hand sequences of all individuals were first concatenated to increase coverage. We designated an acceptable Phred-score of 20 and a 95% cutoff on reads with bases less than this score. This was followed by trimming reads to a uniform 100 bp to allow straightforward calculation of coverage percent downstream. Reads shorter than 100 bp were discarded. Left- and right-hand reads were then interlaced and scanned for overlap. Next, we used RepeatExplorer v2.3.8, which uses a graphical clustering approach to identify repeat profiles using sequencing reads and has been shown to perform well with low-coverage sequencing (Novák et al 2010; Lower et al 2017; Hloušková et al 2019). We specified the Metazoa 3.0 protein database for repeat annotation. For each population, we ran RepeatExplorer for the maximum number of processible reads, which ensured coverage was ≥ 1% for all populations (**Table 1**). Only clusters that occurred in > 0.01% of the sampled reads were annotated, the remaining fell into a “bottom clusters” category.

### Phylogenetic inference and comparative methods

To determine whether populations of the same “species” could be considered as independent evolutionary lineages (i.e., represented cryptic diversity), we performed pairwise genetic differentiation tests in DnaSP v6 (Rosas et al 2017) with 5–10 individuals per population using the mitochondrial locus. We performed the three measures of differentiation from Hudson et al (1992), namely *H*_ST_, *K*_ST_, and *K*_ST_*. The first, *H*_ST_, is measured as 1 – (*H*_S_ / *H*_T_), where *H*_S_ is the weighted average of haplotype diversity within the subpopulation and *H*_T_ is the total population haplotype diversity. *K*_ST_ is measured as 1 – (π_12_ / π), where π_12_ is the weighted average of nucleotide differences between site 1 and 2, and π is the average number of differences irrespective of locality. Finally, *K*_ST_* is identical to *K*_ST_ except π_12_ is changed to log(1 + π_12_), which acts to downweight π_12_ when it becomes high (Hudson et al 1992). Significance was determined using a permutation test with 1000 replicates. Populations with significant differentiation values (α = 0.05) in at least two of the three tests were retained as “independent”; otherwise, one sample site was randomly discarded.

To estimate the age of the clade and to perform phylogenetic comparative methods, we first constructed a time-calibrated tree using BEAST2 (Bouckaert et al 2019). We specified a GTR model of substitution for *COI* and an HKY model for *28S*, as these were the indicated best models in PartitionFinder2 (Lanfear et al. 2017). We applied a molecular clock to *COI* as an exponential prior with a mean of 0.01 substitutions per site per Myr (Hancock et al 2019; Takada et al 2018; Knowlton & Weigt 1998). We applied a relaxed lognormal clock to *28S* following Hancock et al (2019). We then applied two calibration points based on known vicariant events: 1) the closure of the Okefenokee Trough ~1.75 Myr separating Atlantic and Gulf taxa (Bert 1986; McClure & Greenbaum 1999); and 2) the proposed interglacial Pleistocene colonization of *H. arenarius* to Europe (Bousfield 1970). For the first calibration, we applied an exponential prior offset to 1.5 Myr (a conservative lower bound for the closure of the trough) to the *Acanthohaustorius sp*. and *A. millsi* clade, GoM and Atlantic endemics, respectively. The second calibration we applied a gamma distributed prior with α = 1.5 and *β* = 1.0, which has a mean of ~1.18 Myr, roughly in the middle of the Pleistocene. The MCMC was run for 50 million generations ensuring ESS values > 200. We generated a consensus tree using TreeAnnotator (Bouckaert et al 2019), which was visualized in FigTree v1.4.3 (Rambaut 2011).

The resulting tree was imported into the R platform and pruned down to only the taxa in which genome size data was available using the package *ape* v5.3 (Paradis & Schliep 2019). We next used the package *caper* v1.0.1 (Orme et al 2018) to perform phylogenetic generalized least squares (PGLS) on associations between genome size and environmental variables, body size, and repeat element profiles. This allowed us to estimate Pagel’s λ, which ranges from 0.0–1.0 and indicates how much phylogenetic signal is present in the data (Pagel 1999). In addition, we performed ancestral state reconstructions on both the continuous and discrete variables using the package *phytools* (Revell 2012).

## Results

### Genome size variation and age of the clade

Estimates from flow cytometry indicated genome sizes ranged from 2130 Mb to 13080 Mb (**Table 1**). The amphipods with the smallest genomes were all restricted to warm, brackish water – namely, *H. allardi* and the *L. triarticulatus* species complex. The largest genomes belonged to the *H. canadensis* complex, with the largest (13080 Mb) found at Tybee Island, Georgia.

All pairwise differentiation comparisons were significant for at least two measures except for between *H. jayneae* sampled at Carrabelle Beach, FL and Grayton Beach, FL / Dauphin Island, AL (**Table S1**). Carrabelle Beach was thus dropped from subsequent analyses.

The calibrated phylogenetic tree indicated that the entire haustoriid clade is less than 12 Myr old, and that the clade in which we performed genome size estimates is only ~3.5 Myr old (**Figure 1c**); however, the range of the 95% highest posterior density (HPD) was large indicating wide uncertainty in the age estimates (1.67–6.25 Myr). The estimated rate for *COI* was 0.044, which is ~4x faster than previous clock estimates for this gene (Knowlton & Weigt 1998).

Examination of the log files indicated that the calibration prior on the *Acanthohaustorius* clade strongly limited the MCMC sampling as the posterior distribution pressed against the 1.5 Myr lower-bound (**Figure S1**). This indicates that the data may support a younger age for the *Acanthohaustorius* split than the vicariant event will allow. Alternatively, the calibration prior for the *H. canadensis–H. arenarius* split departed from the 1.18 Myr mean rapidly, and the 95% HPD settled around a median of 2.65 Myr, indicating the data support an age for this split that is deeper than the Pleistocene.

### Repeat profiles

The most abundant annotated repeat elements were Class I repeats, the genomic proportion of which ranged from 3.9% (*L. triarticulatus*, Pensacola Bay, FL) to 48.6% (*Neohaustorius schmitzi*). Of the Class I elements, LINEs were the most abundant, accounting for as much as 29% of the genome (*H. jayneae*, Dauphin Island, AL). LTRs were less prevalent, normally less than 6% (see **Table S2**), but two taxa showed dramatic LTR expansions: *N. schmitzi* (18.9%) and *L. dysticus* (17.7%). The only Class II repetitive element identified was the large transposon *Maverick*, which made up at most 1.5% of the genome (*L. dysticus*). Large satellite repeats were identified in all species, which ranged from 1.5% (*H. canadensis*, Melbourne, FL) to 28.5% (*L. triarticulatus*, Pensacola Bay, FL) of the genome. Unlike the previous repeat elements, satellite repeats appeared to be the most prevalent in the smallest genomes (**Figure 2a**; **Table S2**).

**Figure 2.**
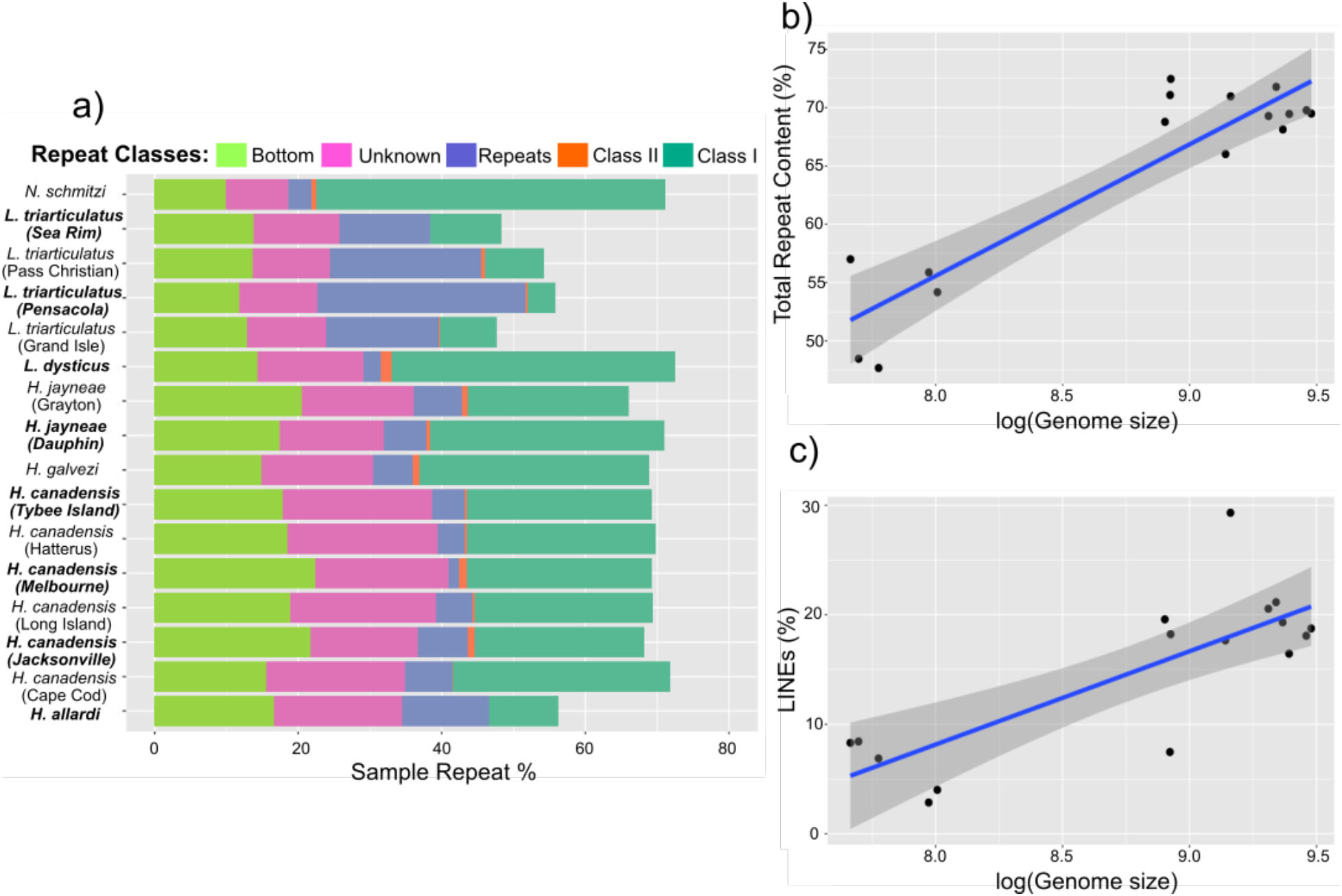
Repetitive content. a) Barplot of repeat content proportion per sample, colored by repeat element family, every other species name bolded to aid in visualization; b) correlation between log(genome size) and total repetitive content; c) correlation between log(genome size) and LINE proportion.

### Genome size correlates and ancestral state reconstructions

We found no correlation between latitude and genome size with either the raw data (*p* = 0.08, *r*^2^ = 0.19) or incorporating phylogeny (*p* = 0.20, *r*^2^ = 0.11). We also found no correlation between genome size and salinity (*p* = 0.67, *r*^2^ = 0.01). However, we found a strong correlation between body length and genome size in both the raw (*p* = 0.01, *r*^2^ = 0.33) and the PGLS comparisons (*p* = 0.01, *r*^2^ = 0.33). The maximum-likelihood (ML) estimate for Pagel’s *λ* = 0.0, indicating there was no phylogenetic signal in body length. For the body length mixture models, we found the model with the lowest AIC score (60.18) was one that included genome size (*p* = 0.01), temperature (*p* = 0.04), and an interaction between genome size and temperature (*p* = 0.02; see **Table 2**). In addition, we found a strong positive correlation between latitude and body size (*p* = 0.0007, *r*^2^ = 0.57, *λ* = 0.0).

**Table 2.** Summary of genomic correlation models. Abbreviations: TR (total repeat %), GS (genome size), C1 (Class I repeats), C2 (Class II repeats), RE (satellite repeats), LTR (LTR-retrotransposons), LI (LINEs), BO (bottom clusters), Lat (latitude), BL (body length), SA (salinity), T (temperature).

| Model | df | *P* value | *r*<sup>2</sup> | Pagel's $\lambda$ | $\lambda$ 95% CI | AIC |
| --- | --- | --- | --- | --- | --- | --- |
| TR ~ log(GS) | 14 | **4.713e-05** | 0.7052 | 0.824 | 0.127–0.961 | - |
| C1 ~ log(GS) | 14 | **0.004834** | 0.4439 | 0.872 | 0.481–0.970 | - |
| C2 ~ log(GS) | 14 | 0.1064 | 0.1754 | 0.000 | 0.000–0.813 | - |
| RE ~ log(GS) | 14 | **0.04179** | 0.2639 | 0.907 | 0.000–0.994 | - |
| LTR ~ log(GS) | 14 | 0.194 | 0.1174 | 0.995 | 0.938–1.000 | - |
| LI ~ log(GS) | 14 | **0.001314** | 0.5334 | 0.491 | 0.000–0.856 | - |
| BO ~ log(GS) | 14 | 0.1513 | 0.1413 | 0.514 | 0.000–0.914 | - |
| log(GS) ~ log(Lat) | 14 | 0.2013 | 0.1138 | 1.000 | 0.981–1.000 | - |
| BL ~ Lat | 14 | **0.0007267** | 0.5696 | 0.000 | 0.000–0.827 | - |
| log(GS) ~ SA | 14 | 0.6787 | 0.01262 | 1.000 | 0.958–1.000 | - |
| BL ~ T + log(GS) +<br>T*log(GS) | 12 | **0.0001575** | 0.8027 | 0.000 | 0.000–0.870 | **60.17677** |
| BL ~ T + log(GS) | 13 | **0.0004119** | 0.6986 | 0.000 | 0.000–0.824 | 64.96170 |
| BL ~ T | 14 | **0.0007938** | 0.5644 | 0.490 | 0.000–0.945 | 67.21072 |
| BL ~ log(GS) | 14 | **0.01929** | 0.3329 | 0.000 | 0.000–0.744 | 75.67192 |

We found a positive relationship between genome size and total repetitive content (*p* = 0.000047, *r*^2^ = 0.71), and a ML estimate of Pagel’s *λ* = 0.82, indicating that there was strong phylogenetic signal in total repeat content (**Figure 2b**). Of the different identified repeat families, Class I repeats had the strongest correlation with genome size (*p* = 0.004, *r*^2^ = 0.44, *λ* = 0.87), with this pattern being driven by LINEs (*p* = 0.001, *r*^2^ = 0.53, *λ* = 0.49; **Figure 2c**). An opposite relationship was found with satellite repeats due to the preponderance of these in the smallest genomes (*p* = 0.04, *r*^2^ = 0.26, *λ* = 0.91). There was no relationship between percent coverage and total genomic repeat content (*p* = 0.34, *r*^2^ = 0.06).

The ancestral state reconstruction analysis revealed a general correlation between increases in genome size and body length, with the latter lagging behind the former (**Figure S2**). In addition, using the full haustoriid phylogeny we find that the ancestral environment was most likely an open coast with high salinity, and that there have been at least four independent transitions into brackish, protected beaches: 1) *Eohaustorius* on the Pacific coast; 2) the *L. triarticulatus* species complex in the GoM; 3) *L. dysticus* in the North Atlantic; 4) and *H. allardi* along the Texas and Louisiana coastline (**Figure 3**).

**Figure 3.**
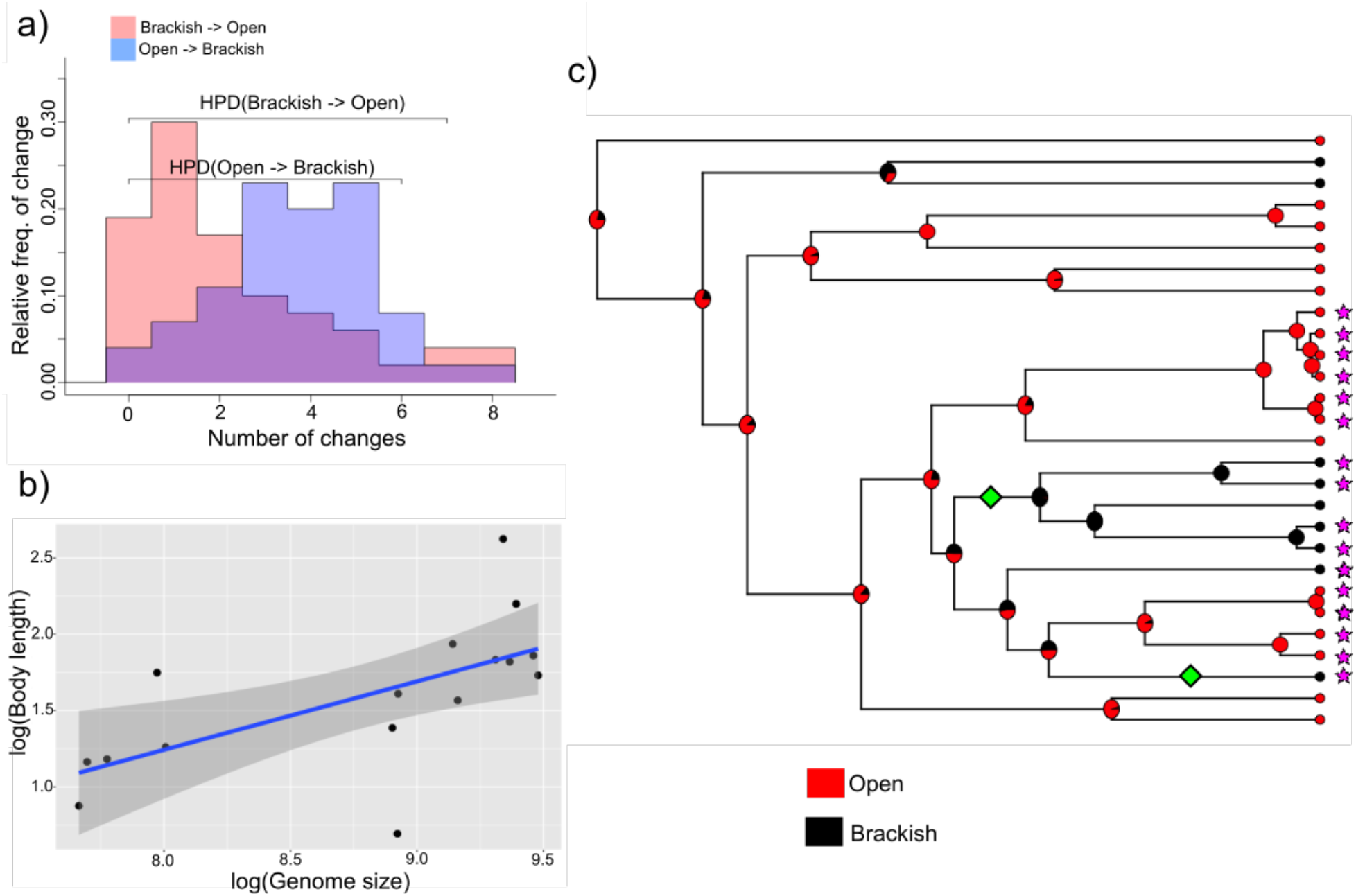
Transitions to brackish water. a) transition densities between brackish and open waters; b) correlation between log(genome size) and body length; c) ML estimate of ancestral environment type (“brackish”, “open”). Red is “open”, black is “brackish”. Green diamonds represent transitions to warm brackish bays and lineages with genome size reductions; purple stars are samples with genome size estimates.

## Discussion

Is there a general “genome-temperature-size-rule” that explains haustoriid genome evolution? We find mixed evidence for this, indicating that the factors that influence genome size are complex and multifaceted. While we find evidence from mixed models of an interaction between genome size and temperature on body length, there was not a significant relationship between latitude and genome size (*p* = 0.08). Instead, genome size variation in haustoriid amphipods may be the result of differing environments, with amphipods living on surf-swept, high salinity beaches having increased body length and genome sizes compared to those living in warm, brackish bays. Further, we find that genome size and repeat element prevalence have strong phylogenetic signal (**Table 2**).

### The drivers of genome size evolution in Haustoriidae

Haustoriid amphipods display incredible genome size variation across relatively recent evolutionary time. Most haustoriid genomes are relatively large (>7000 Mb), but this may be an ancestral condition. The most closely related amphipod in the Animal Genome Size Database (http://www.genomesize.com/; Gregory 2020) are the pontopoeriids *Monoporeia affinis* and *Pontoporeia femorata*, each of which have genome sizes of 8242 Mb. Therefore, *N. schmitzi, L. dysticus, H. galvezi*, and *H. jayneae* may represent cases of relative genomic stasis (± 1000 Mb). Alternatively, the *H. canadensis* clade has experienced rapid genomic expansions within the last 3 Myr. It should be noted that *H. canadensis* and its sister species, *H. arenarius*, are some of the largest fossorial amphipods and are the most abundant at higher latitudes.

In stark contrast, the predominantly warm, brackish water clades of *L. triarticulatus* and *H. allardi* show evidence of massive genomic purging. If the ancestral genome size was ~8000 Mb, these clades have lost on average 5500 Mb. These genomes are largely devoid of LINEs, which occupy up to 30% of the genomes of their relatives. This may indicate that genomic purging has targeted transposable elements specifically, which has been found in other cases of genomic purging (Kelley et al 2014; Barrón et al 2014; Lyu et al 2018; Misof 2019). Notably, these genomes have a much higher percentage of satellite repeats than their relatives, especially in the *L. triarticulatus* species complex. Since these repeats are distributed throughout this complex, it likely represents an ancestral expansion.

Brackish bays present unique physiological stresses not present on the open coast. Bays tend to be on average warmer than the open ocean, and in the GoM can reach 34°C in the summer months (Ross & Behringer 2019; Holmquist et al 1989; this study). These higher temperatures reduce the capacity of water to retain dissolved oxygen, which increases respiratory stress compared to the highly oxygenated sands of the open coast (Bousfield 1970). Bays also have shallow anoxic layers due to densely packed organic matter (Revsbech et al 1980). Finally, due to freshwater runoff and poor mixing, bays are typically brackish and may have salinities as low as 5‰ (this study), which increases osmotic stress on cells. Haustoriid amphipods are in general tolerant of wide salinity fluctuations, but only a few clades have specialized to live in bays and freshwater inlets; namely, *Eohaustorius estuarius*, the *L. triarticulatus* species complex, *L. dysticus*, and *H. allardi*. These species may also be found on the open coasts, but they tend to be less numerous and relegated to the subtidal zone (LeCroy 2002; Hancock & Wicksten 2018).

Temperature and salinity have been shown to have synergistic impacts on oxidative stress, such that species may tolerate high temperatures or low salinity, but not in conjunction (Takolander et al 2017). This may explain why the two GoM shifts to brackish environments are accompanied by body size and genome size reductions, but not the Atlantic shift (i.e., *L. dysticus*; **Figure 3c**). The North Atlantic consistently has lower sea surface temperatures (SST) than the GoM, and therefore for much of the range of *L. dysticus* it may be exposed to low salinity but not high temperature.

Given the increased metabolic stress associated with transitioning to warm brackish bays, we hypothesize that strong selection on faster metabolism led to rapid genomic purging in *L. triarticulatus* and *H. allardi*. In contrast, reduced metabolic stress at higher latitudes in *H. canadensis* may have provided a selectively permissive environment for transposable elements to proliferate.

Unfortunately, we were not able to distinguish within-complex trends in genome size evolution due to low sample sizes. Genome sizes were generally more similar the more phylogenetically related two samples within a complex were, but without additional material we cannot conclude if this is diagnostic or a random sampling effect.

### Body size evolution in Haustoriidae

Haustoriid amphipods range from 2–18 mm in length from rostrum to telson, and even within a single species complex may range from 4–18 mm (i.e., *H. canadensis*). Three major patterns of body size evolution emerged from this study: 1) influence of temperature and latitude; 2) influence of sand grain size; and 3) influence of genome size.

Previous work has established a relationship between body length and latitude (e.g., LeCroy 2002; Hancock & Wicksten 2018), and this extends beyond haustoriid amphipods (Hultgren et al 2018). We recapitulate these results, finding a strong correlation between body length and latitude. Interestingly, despite this correlation and the support from the mixed model of an interaction between genome size and temperature on body length, genome size is not correlated with latitude (*p* = 0.08). This discrepancy may be the result of temperature variation at individual sites irrespective of latitude. This is related to the “bay versus open ocean” discussion above, in which the topography of a given sample site may influence its temperature profile. This helps to explain why there is a similar disparity in body length across the GoM, despite little differences in overall SST averages. However, the higher summer temperatures undoubtedly constrain body sizes in the GoM, as not even the largest sampled haustoriid approaches the size sampled at the northernmost latitudes (9 mm versus 18 mm).

Crustacean body sizes are highly plastic and often environmentally determined (Cheng & Chang 1994; Atkinson & Sibly 1997; Twombly & Tisch 2000). Hancock & Wicksten (2018) posited that variation in sand grain size at different sites may help explain body size differences within a single species range. Fine sand predominates in the western GoM and tends to be more compact, which would be difficult to move through for a large amphipod (**Table 2**). Alternatively, in the eastern GoM and Atlantic, coarse and medium sand is the most common. Coarse sand is loose and characterizes beaches with steep berms and strong surf. These factors together may promote larger body size to traverse the heavy quartz sand and reduce the chance of being dislodged from the sediment by wave action.

Outside of latitude, genome size was the strongest predictor of body size. Genome sizes are known to increase cell size (Gregory 2005), but this does not necessarily lead to a proportional increase in body size. Instead, the relationship between body size and genome size may be an emergent property of the previous two factors impacting body size, and, perhaps most importantly, the strength of selection on metabolism.

### Conclusions

Genome size variation is likely the result of a complex network of interactions that include cellular and metabolic processes, life-history, and the population genetic environment (Gregory 2005; Lynch 2007). We have shown that that variation can arise rapidly following shifts to new environments, whether to a more selectively permissive or constraining one. Future work on haustoriid amphipods should include more comprehensive genome size sampling across the phylogenetic tree, especially in other species that have made the transition to brackish waters (such as *E. estuarius*). In addition, laboratory studies examining the physiological tolerances of different haustoriids would be valuable to test the hypotheses proposed here.

## Acknowledgements

We thank the many people who housed us as we collected samples for this work, especially Liz Marchio and Joël Henrico. This work was funded by Texas Sea Grant.

