## Supplementary Material for "Rapid genomic expansion and purging associated with habitat transitions in a clade of beach crustaceans (Haustoriidae: Amphipoda)"

**Tables**

Table S1. Pairwise genetic differentiation measures.

| **Comparison** | ***H*_ST_** | ***K*_ST_** | ***K*_ST_*** |
| --- | --- | --- | --- |
| *H. jayneae* (Dauphin Island) x *H.* *jayneae* (Grayton Beach) | **0.21032*** | **0.54839*** | **0.42426*** |
| *H. jayneae* (Dauphin Island) x *H. jayneae* (Carrabelle Beach) | 0.12348 | 0.27273 | 0.20226 |
| *H. jayneae* (Grayton Beach) x *H. jayneae* (Carrabelle Beach) | 0.04545 | 0.06604 | 0.2850 |
| *H. canadensis* (Cape Cod) x *H. canadensis* (Long Island) | **0.20687**** | 0.15385 | **0.15438*** |
| *H. canadensis* (Cape Cod) x *H. canadensis* (Melbourne) | **0.1340*** | **0.72846***** | **0.44586***** |
| *H. canadensis* (Cape Cod) x *H. canadensis* (Jacksonville) | **0.59241**** | **0.83879**** | **0.70653**** |
| *H. canadensis* (Cape Cod) x *H. canadensis* (Tybee) | **0.15020**** | **0.74960***** | **0.44608**** |
| *H. canadensis* (Long Island) x *H. canadensis* (Jacksonville) | **0.64770**** | **0.87097**** | **0.80899**** |
| *H. canadensis* (Long Island) x *H. canadensis* (Melbourne) | 0.04698 | **0.76483**** | **0.46874**** |
| *H. canadensis* (Long Island) x *H. canadensis* (Tybee) | 0.04698 | **0.74251**** | **0.41554**** |
| *H. canadensis* (Jacksonville) x *H. canadensis* (Melbourne | **0.462628***** | **0.4662***** | **0.482366***** |
| *H. canadensis* (Melbourne) x *H. canadensis* (Tybee) | 0.01296 | **0.4400**** | **0.34241***** |

Table S2. Proportion of major clusters.

| **Species** | **Genome size (Mb)** | **Max reads** | **Coverage (%)** | **Total Repeats (%)** | **LINE** | **DIRS** | **LTR** | **Penelope** | **Maverick** | **Satellite** | **Unknown** | **Bottom** |
| --- | --- | --- | --- | --- | --- | --- | --- | --- | --- | --- | --- | --- |
| *Nschmitzi* | 7500 | 1655500 | 0.0221 | 71.08 | 7.47 | 7.09 | 18.86 | 0.1 | 0.6 | 2.78 | 8.78 | 9.83 |
| *HcanadensisTy* | 13080 | 2145387 | 0.0164 | 69.49 | 18.75 | 0 | 3.74 | 0.4 | 0.28 | 4.63 | 20.81 | 17.81 |
| *HjayneaeG* | 9330 | 2464505 | 0.0264 | 66.02 | 17.66 | 0.22 | 3.31 | 0.51 | 0.77 | 6.81 | 15.56 | 20.47 |
| *HjayneaeDI* | 9520 | 1420106 | 0.0149 | 70.97 | 29.31 | 0.01 | 2.09 | 0.09 | 0.61 | 5.79 | 14.49 | 17.38 |
| *Hallardi* | 2130 | 1128376 | 0.053 | 57 | 8.31 | 0.01 | 0.98 | 0.32 | 0.31 | 11.31 | 17.96 | 16.57 |
| *HcanadensisCC* | 11390 | 1845132 | 0.0162 | 71.78 | 21.15 | 1.52 | 4.44 | 0.21 | 0.06 | 6.39 | 19.3 | 15.58 |
| *HcanadensisNC* | 12830 | 2469903 | 0.0193 | 69.77 | 18.08 | 0.54 | 5.11 | 0.47 | 0.31 | 3.6 | 21.02 | 18.47 |
| *HcanadensisLo* | 11990 | 2585215 | 0.0216 | 69.46 | 16.45 | 0.57 | 5.51 | 0.39 | 0.33 | 4.93 | 20.27 | 18.92 |
| *LdysticusNCBa* | 7520 | 1857818 | 0.0247 | 72.46 | 18.22 | 3.14 | 17.65 | 0.49 | 1.47 | 2.02 | 14.84 | 14.28 |
| *HcanadensisJa* | 11690 | 2889172 | 0.0247 | 68.13 | 19.31 | 0 | 3.94 | 0.36 | 0.86 | 6.69 | 14.92 | 21.72 |
| *HcanadensisMe* | 11050 | 2354941 | 0.0213 | 69.28 | 20.55 | 0.38 | 4.42 | 0.43 | 1.13 | 1.47 | 18.51 | 22.41 |
| *HgalveziMX* | 7350 | 2501302 | 0.034 | 68.78 | 19.58 | 3.93 | 6.33 | 1.6 | 0.95 | 5.6 | 15.53 | 14.84 |
| *LtriGI* | 2380 | 644675 | 0.0271 | 47.67 | 6.88 | 0 | 0.94 | 0.1 | 0.13 | 15.28 | 10.98 | 12.89 |
| *LtriS* | 2200 | 785415 | 0.0357 | 48.46 | 8.43 | 0.04 | 1.44 | 0.07 | 0.16 | 12.56 | 11.92 | 13.84 |
| *LtriPC1* | 3000 | 316053 | 0.0105 | 54.17 | 4.01 | 0 | 4.1 | 0.04 | 0.57 | 20.8 | 10.85 | 13.58 |
| *LtriPBAY* | 2900 | 268152 | 0.0092 | 55.87 | 2.86 | 0 | 1.06 | 0.01 | 0.25 | 28.51 | 10.82 | 11.77 |

**Figures**

**Figure S1.** Posterior estimate of the age of the *Acanthohaustorius* clade.

**Figure S2**. Ancestral reconstruction of genome size against body size.

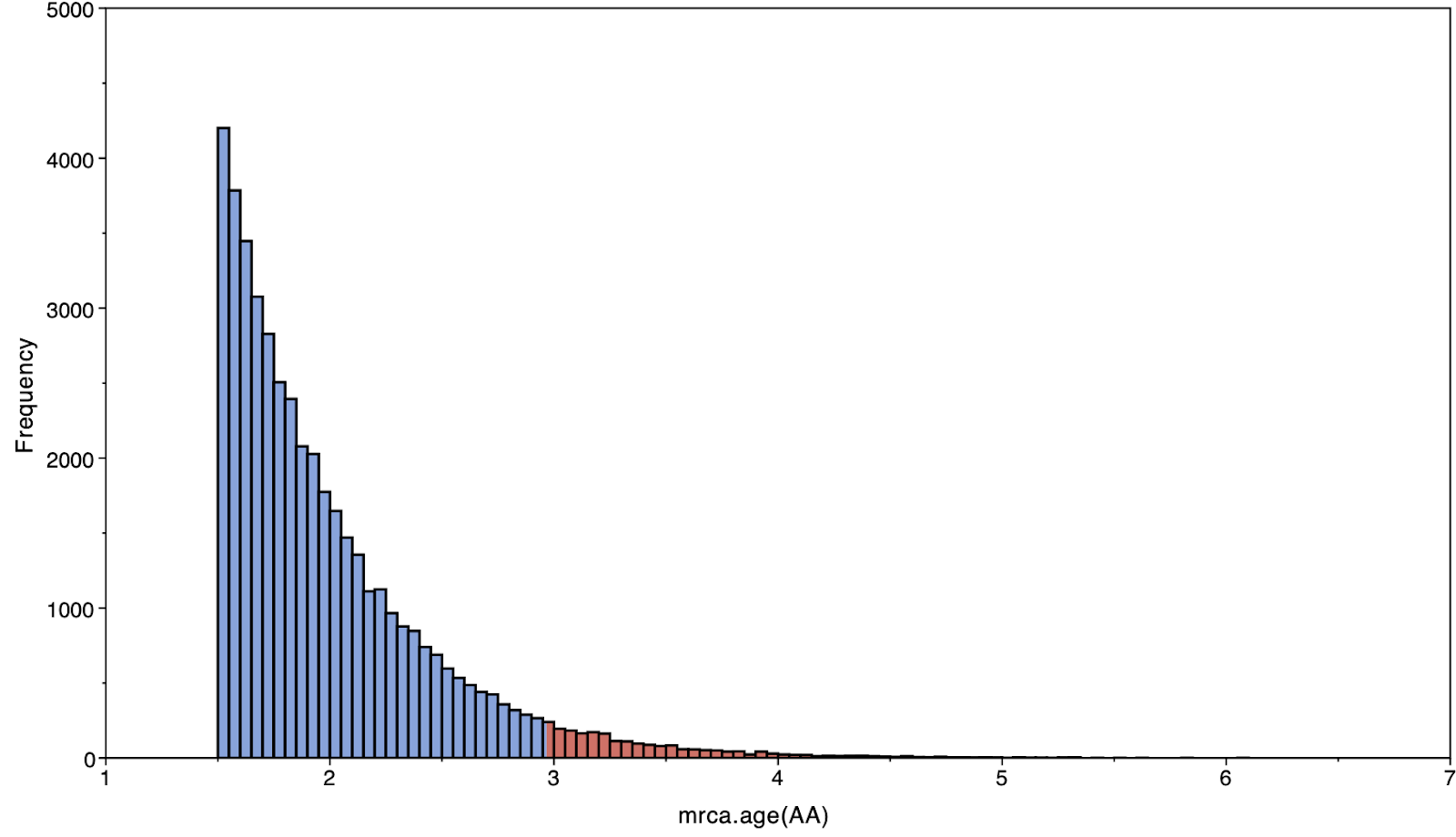

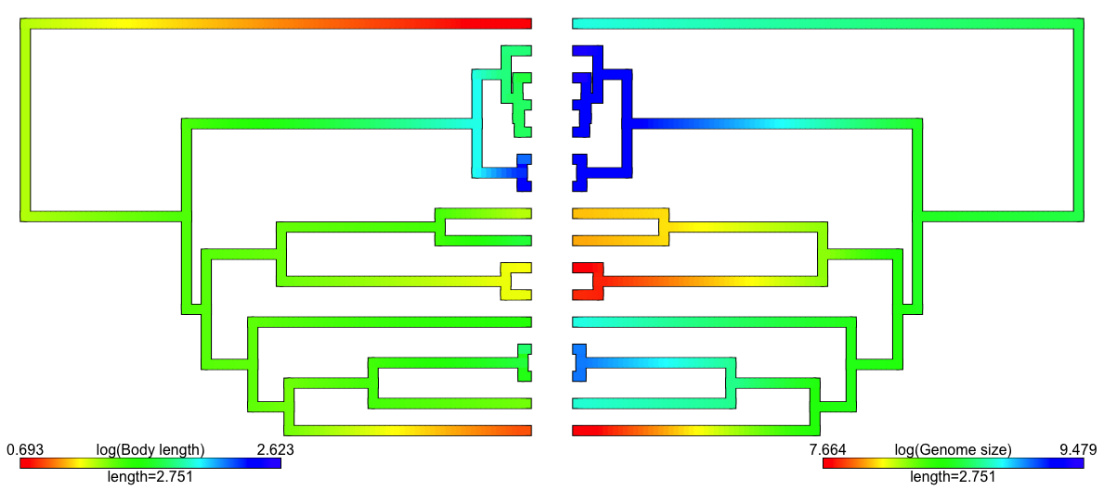
